# *TailTimer:* an open-source device for automating the rodent tail immersion assay

**DOI:** 10.1101/2020.09.25.313858

**Authors:** Mallory E. Udell, Angel Garcia Martinez, Tengfei Wang, Jie Ni, Christian Hurt, Megan Mulligan, Eva E. Redei, Hao Chen

## Abstract

The tail immersion assay is a widely used method for measuring acute thermal pain in a way which is quantifiable and reproducible. It is non-invasive and measures response to a stimulus that may be encountered by an animal in its natural environment. However, tail withdrawal latency data are usually collected manually, and precise temperatures of the water at the time of measurement are most often not recorded. These two factors can reduce the reproducibility of tail immersion assay data. We designed a device, *TailTimer*, which uses the Raspberry Pi single-board computer and a temperature sensor, to automatically record both tail withdrawal latency and water temperature. The device has a radio frequency identification (RFID) system that can record the ID of animals. Our software recognizes several specific RFID keys as user interface commands, which allows *TailTimer* to be operated via RFID fobs. We also programmed the device to only allow tests to be conducted when the water is within ± 0.25 °C of the target temperature. Data recorded using the *TailTimer* device showed a linear relationship between tail withdrawal latency and water temperature when tested between 47 - 50 °C. We also observed a profound effect of water mixing speed on tail withdrawal latency. Our data further revealed significant strain and sex differences, valorizing *TailTimer* in its ability to detect genetically-determined variations in thermal pain sensitivity.

**Significance Statement:** Quantification of tail withdrawal latency in response to thermal pain has essentially remained the same since the method was first introduced decades ago and relies on manual recording of water temperature and tail withdrawal latency. Such manual methods engender relatively substantial variability and are potential contributors to some of the discrepancies present among relevant research. The open source *TailTimer* device we report here is simple and inexpensive to manufacture. The RFID-based user interface is ergonomic, especially in animal facilities where space is limited and gloves are mandatory. We anticipate that the increased reproducibility of tail withdrawal latency provided by *TailTimer* will augment its utility in nociception and addiction research.

## Introduction

Extant research has utilized an assortment of methods to measure and understand nociception relative to the specific type of pain being assessed. Both acute and chronic pain can be modeled in rodents. Moreover, pain can be induced through the use of various types of stimuli including thermal, electrical, chemical, and mechanical. In order to properly assess acute pain, it is imperative that the noxious stimuli meets the requirements of being quantifiable, reproducible, and non-invasive. Furthermore, the stimulation must be delivered promptly and briefly enough to induce synchronous excitation of the nerve fibers (Le Bars et al., 2001). For this reason, neither mechanical nor chemical stimulation are adequate for studying acute pain. Electrical stimulation, though specific, is disadvantageous in that it does not reflect a type of stimuli that an animal would encounter in its natural environment. The use of thermal stimulation is therefore conducive for modeling acute pain, as heat is more selective and naturally encountered (Le Bars et al., 2001).

The tail immersion assay is a thermal assessment of pain that is often used to determine the analgesic properties of drugs. The assay measures pain by immersing a rat’s tail in hot water and recording the time it takes before a tail flick (i.e., pain response) is observed. However, current standards for data collection often rely on manual recording and monitoring, inviting an increased likelihood for error. The use of manual methods may thereby contribute to some of the inconsistent findings among previous pain studies (Fischer et al., 2008; Picker et al., 2011). Thus, the development of a more reliable method for this simple assay is warranted.

The current study describes the open source *TailTimer* device as an automated method for measuring acute thermal pain using the tail immersion assay. This device uses a single-board computer (i.e., Raspberry Pi 3, RP3), which offers an affordable alternative to desktop computers without compromising capability. In addition, RP3 can be paired with a variety of devices and sensors to offer controllability of a wide range of internal and environmental variables. Here, we added a digital temperature probe and a radiofrequency identification (RFID) system. This automated method abates the burden of data recording while, more importantly, ensuring precise measurement of tail withdrawal latencies and water temperature.

## Materials and Methods

### Hardware

The *TailTimer* is composed of a Raspberry Pi 3 (Model B, RaspberryPi Foundation, UK) single-board computer (RP3), a 5-inch touch screen (DFR0550, DFRobot, ShangHai, China), a waterproof digital temperature sensor (DS18B20, Adafruit Industries, NY, USA), an RFID reader (EM4100, 125 kHz, HiTag, available at Amazon.com), and two electrical wires. The ground wire is connected to one of the ground pins of the general purpose input/output (GPIO), and the latency wire is connected to GPIO 18 (pin 12). All components are enclosed in a 3D-printed case (see Fig. S1). The connection of the components is outlined in Fig. 1. The temperature sensor connects to the RP3 via a one-wire serial interface to provide continuous temperature readings. The RFID reader connects to the RP3 via a USB port. It scans an RFID chip embedded under the skin of a rat. Each RFID chip contains a unique code that serves as the ID of a rat. During operation, the ground wire and thermal probe remain immersed together in the hot water at approximately 50% of the depth. The latency wire is dipped into the hot water at the same time as a rat’s tail and is taken out of the hot water, together with the tail, when a pain response is observed (i.e., the tail starts to “flick” in response to heat).

**Figure 1.**
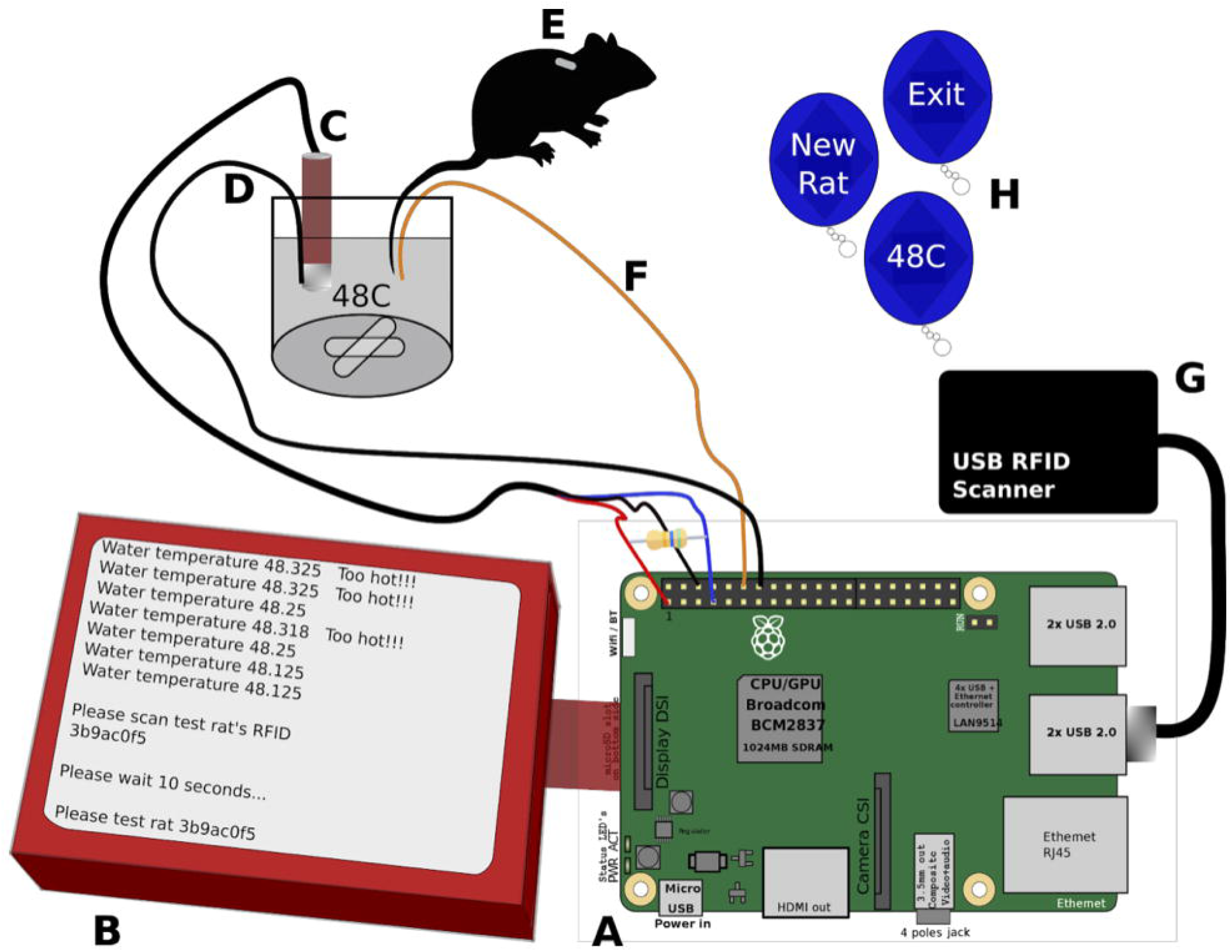
Components of the *TailTimer* device for use in the tail immersion assay. **(A)** The RP3 computer. (B) The *TailTimer* software in operation. (C) Thermal probe to detect water temperature. (D) Electrical ground wire that remains immersed in the water with the thermal probe (C). **(E)** Scannable RFID implanted subcutaneously in the rat for identification. **(F)** Electrical latency wire to be dipped into and withdrawn from the water simultaneously with the rat’s tail to start and stop the timer, respectively. **(G)** USB RFID scanner. **(H)** Scannable RFID command fobs used to navigate the *TailTimer* program in place of a keyboard and mouse. The six necessary fobs are used to enter the user ID, set the target temperature, start a new rat, test the same rat again, delete the last latency, and exit the program.

### Hot water

A standard hot plate (Thermolyne Cimarec 1, Model SP46615) was used to heat a 1 L beaker containing 900 ml tap water. Water temperature was adjusted to 48 ± 0.25 °C. To ensure homogeneity of the temperature, a 2 cm magnetic stir bar was used to mix the water at a consistent, low speed (setting one on the hot plate). Approximately 9 g of NaCl (1%) was added to increase conductivity of the water, as required for automated recording of the open/close state between the ground and latency wires.

### Software features, user interface, and testing procedure

*TailTimer* has a computer program written in the Python language. It collects information from the temperature probe and the RFID reader and logs the number of seconds during which the circuit between the latency wire and the ground wire is in the close state (i.e., tail withdrawal latency). A text-based user interface is automatically started when the device is powered on. It first prompts the user to input a username, set the target temperature, and test the connection of the wires. Measures of tail withdrawal latency are then initiated by entering the ID of the rat. This can be achieved by scanning the RFID embedded under the skin of the rat using the USB RFID reader or by entering the ID via a keyboard. The interface then prompts the user to dip the rat’s tail, together with the latency wire, into the hot water at a depth of approximately 4 cm. A timer starts when the latency wire contacts the water and stops when it is withdrawn. Tails were removed from the water if the rat failed to elicit a response after 10 s. Each rat was tested at least twice. Additional testing was conducted when the difference between a rat’s latencies exceeded 1.5 s. The software enforces a rest period of 10 s in between tests. Assessments can only be performed when the water is within 0.25 °C of the target temperature. The device produces continuous water temperature updates and displays warnings when the temperature falls out of the target range. All relevant data are automatically recorded by the program. The user can operate the program using a keyboard. However, the RFID system offers a more convenient alternative by encoding the limited number of answers to the user interface questions in RFID fobs with each fob representing one unique answer. A total of six fobs (one each for entering the user ID, setting the target temperature, starting a new rat, testing the same rat again, deleting the last latency, and exiting the program) are needed for all operations. Data gathered from the device are transferred using a standard USB drive.

### Software accessibility

The software and design of the 3D enclosure (Fig. S1) described in the paper are freely available in the Github repository at https://github.com/chen42/openbehavior/tree/master/tailTimer.

### Animals

The study was conducted on a total of 45 inbred male and female rats, comprised of three strains: Spontaneously Hypertensive (SHR/NCrl, *n* = 8), Wistar Kyoto more-immobile (WMI, *n* = 15), and Wistar Kyoto less-immobile (WLI, *n* = 22). All rats were naïve to drugs and between 55 and 74 days old at the start of testing. Testing was conducted over the course of two consecutive days. Breeders of the WMI and WLI strains were obtained from Dr. Redei, and the colony was maintained at The University of Tennessee Health Science Center. The rats were housed in groups of two to four without enrichment and maintained in a temperature-controlled room on a 12 h dark/light cycle with lights on at 9:00 pm. Food and water were provided *ad libitum.* This study was conducted in accordance with the NIH Guidelines concerning the Care and Use of Laboratory Animals. All procedures were approved by the Animal Care and Use Committee of The University of Tennessee Health Science Center.

### Measuring tail withdrawal latency in animals under different conditions

Pain assessments were conducted on six WMI males using various water temperatures and mixing speeds. We performed the tail immersion assay on each rat using three different water stirring settings: low speed (setting one), high speed (setting seven), and still (setting turned off) at a temperature of 48 ± 0.25 °C. We also tested these males at adjusted target temperatures of 47, 49, and 50 °C when the water was mixing at a constant low speed (setting one). During testing, the animals were held without restraint. All assessments were performed by the same female human investigator.

### Statistical analyses

Data were presented as means ± SEM. Differences were considered significant at *p* < 0.05. ANOVAs were performed to test for main effects of water temperature with four levels (47, 48, 49, and 50 °C) and water mixing speed with three levels (low, medium, and high) on tail withdrawal latency data. Linear regression analysis using the *R^2^* statistic was conducted to further assess the relationship between water temperature and tail withdrawal latency. We also tested for possible strain-by-sex interactions. Tukey HSD *post hoc* tests were performed to evaluate between-group differences by sex and strain. All analyses were conducted on a laptop computer running macOS High Sierra 10.13.6 using the R-statistical software.

## Results

We designed the *TailTimer* device to quantify tail withdrawal latency automatically by measuring the open time of a circuit between a ground wire that remains immersed in a conductive salt solution and a latency wire that is first dipped into and then withdrawn from the solution together with a tail when the tail starts to “flick” in response to heat. We first calibrated the device’s accuracy of timing by comparing *TailTimer* against two trained technicians. We simulated the assay by dipping the latency wire into and then out of the solution and subsequently compared the latencies recorded using a stopwatch against those recorded by *TailTimer*. We found that *TailTimer* consistently reports approximately 0.4 s shorter latencies than the stopwatch. To account for this, we programmed *TailTimer* to add 0.4 s to each trial.

### Water temperature influences tail withdrawal latency

Six male WMI rats were used to assess tail withdrawal latency at four different water temperatures. Results of a one-way ANOVA revealed a significant main effect of temperature on tail withdrawal latency, *F*_(3,47)_ = 50.93, *p* < 0.001, in which latencies were significantly longer at lower water temperatures and shorter at higher temperatures. Moreover, the difference between latencies measured at 48 versus 49 °C was especially great, with a mean difference of 1.02 s between these two temperatures (*p* < 0.0002). As illustrated in Fig. 2A, results of the linear regression analysis also revealed a significant relationship between water temperature and latency, *F*_(1,49)_ = 156.2, *p* < 0.001, with an *R^2^* of 0.76. The strong linear relationship indicated that the withdrawal latency measured by *TailTimer* is highly accurate.

**Figure 2.**
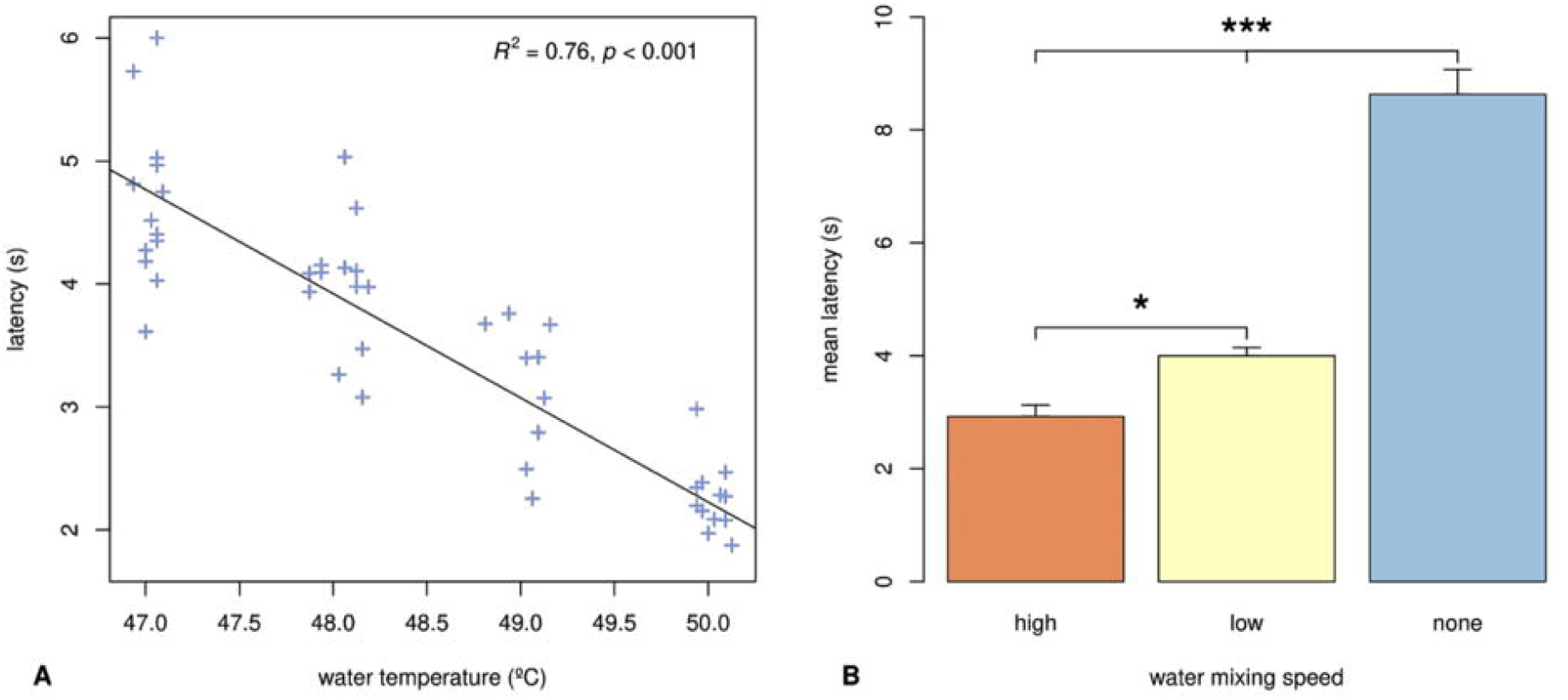
**(A) Tail withdrawal latency measured at different water temperatures.** These findings demonstrate the concomitance between water temperature and latency length. As water temperature is decreased, latency is lengthened. Linear regression indicated that temperature explains 76% of the variance in latency when tested between 47 - 50 °C. **(B) Tail withdrawal latency measured at different water mixing speeds.** Lengthening of latency occurs as the water mixing speed is decreased with the longest latencies occurring when the water is still (i.e., not being mixed by the stir bar). Data are expressed as mean ± SEM; \**p* < 0.05, \*\*\**p* < 0.001.

### Water mixing speed influences tail withdrawal latency

During prior experimentation, we observed an effect of water mixing speed on tail withdrawal latency. Therefore, we used six male WMI rats to systematically measure tail withdrawal latency at 48 °C under three different water mixing speeds. An ANOVA yielded a significant main effect of the water mixing speed on tail withdrawal latency, *F*_(2,35)_ = 110.3, *p* < 0.001. *Post hoc* analysis (Fig. 2B) revealed significantly shorter latencies at the high spin setting (2.94 ± 0.2 s) relative to the low (4.0 ± 0.14 s) and no spin (8.63 ± 0.45 s) conditions. Similarly, latencies were significantly longer when the water was still versus spinning at a low speed (*p* < 0.001). Together, these data elucidate the importance of maintaining consistent mixing speed of the water across experiments. For modified setups, including the utilization of hot water baths, spatial heat gradients should be defined and water temperature should be adjusted accordingly.

### Biological differences in pain sensitivity can be detected by *TailTimer*

We proceeded to use *TailTimer* to detect biological differences (i.e., strain and sex) in thermal pain sensitivity. We measured tail withdrawal latency in three inbred strains of both male and female rats. Results of the two-way ANOVA revealed significant sex differences, *F*_(1,197)_ = 9.517, *p* < 0.01, in which female rats exhibited overall shorter latencies (3.81 ± 0.09 s) relative to their male counterparts (4.27 ± 0.14 s). A significant main effect of strain was also discovered, *F*_(2,197)_ = 149.304, *p* < 0.001. Interestingly, SHR rats displayed significantly longer latencies (6.55 ± 0.29 s) compared to the WLI (3.94 ± 0.07 s) and WMI strains (3.43 ± 0.07 s), which were not significantly different from each other. Most notably, results of the analysis revealed a significant strain-by-sex interaction, *F*_(2,197)_ = 7.582, *p* < 0.001. These results are illustrated in Fig. 3. Significant sex differences were shown within the SHR strain in which latencies were longer in males (7.2 ± 0.32 s) versus females (5.82 ± 0.4 s). Furthermore, *post hoc* results indicated a significant difference between WLI and WMI males (*p* < 0.001) but not between WLI and WMI females. Specifically, we observed significantly longer latencies in WLI males (4.12 ± 0.09 s) versus WMI males (3.4 ± 0.11 s). Together, these findings valorize the *TailTimer* device in its ability to detect genetically-determined differences in thermal pain sensitivity.

**Figure 3.**
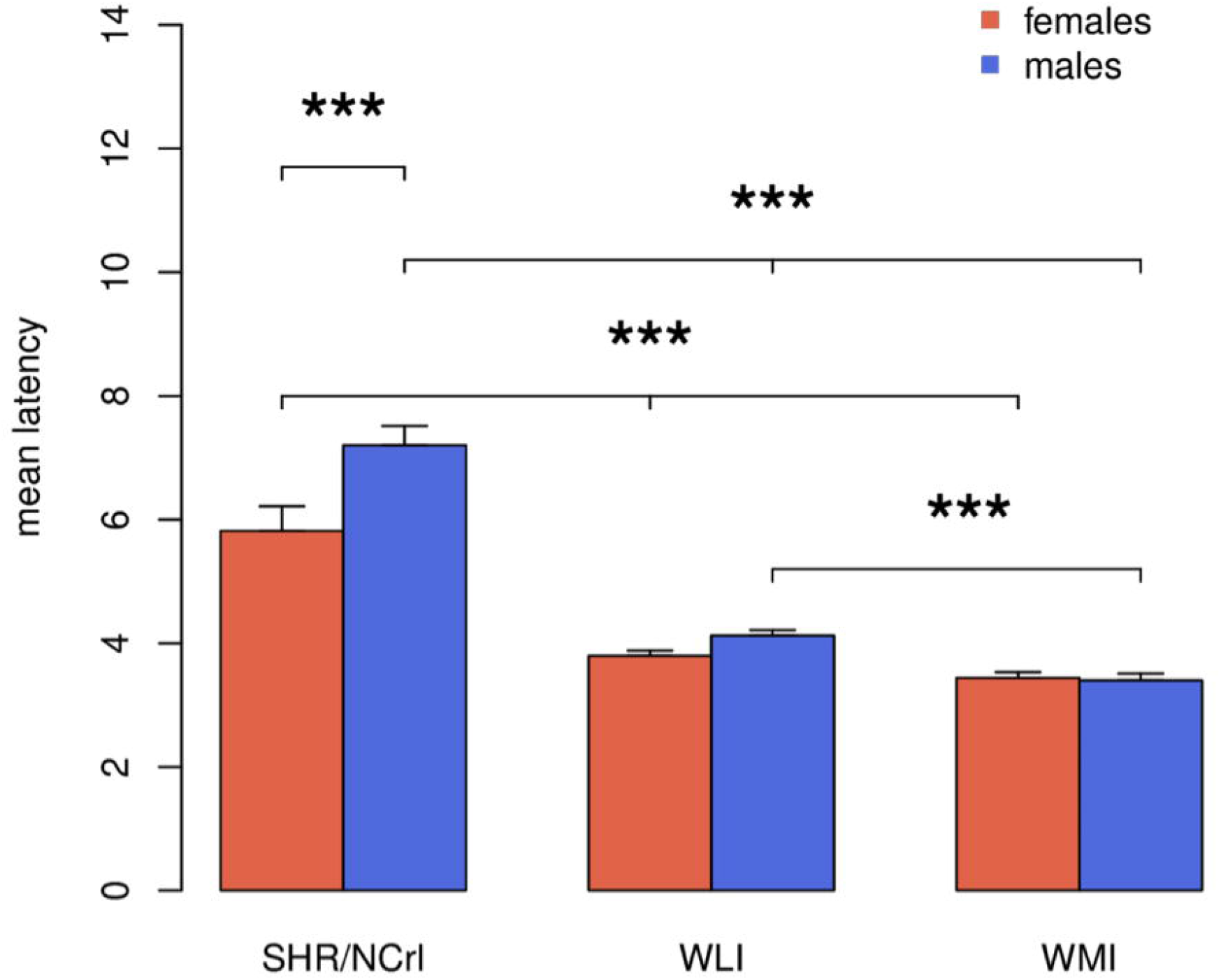
Tail withdrawal latency by sex and strain. Mean latencies reflect the average of the (two - four) tests per individual rat averaged across sex and strain. Data are expressed as mean ± SEM; \*\*\**p* < 0.001.

## Discussion

The purpose of the current study was to develop an automated device for measuring pain sensitivity using the tail immersion assay. Automating data collection and monitoring of variables (e.g., water temperature) abates the burden of data recording while, more importantly, ensuring precise latency measures and water temperature regulation. *TailTimer* records tail withdrawal latency automatically by quantifying the close time of a circuit between a ground wire immersed in a conductive (1%) salt solution and a latency wire that is first dipped into and then withdrawn from the solution, together with a tail, at the first indication of pain (i.e., tail flick). This device is simple to operate and avoids the need to use a stopwatch. It only allows tests to be conducted when water is within ± 0.25 °C of a set target which, in turn, enhances the reproducibility of findings. The technician, however, still needs to observe the movement of the tail carefully and decide when to withdraw the tail and wire from the solution. A single technician should be designated to administer all assessments.

Because tail withdrawal latency is determined by water temperature, we designed the software to only allow tail withdrawal to be measured within a limited range (i.e., ± 0.25 °C). The results of our experimentation conducted on six WMI males using an assortment of target temperatures (47, 48, 49, and 50 °C) further demonstrated the effect of temperature on tail withdrawal latency. As shown in Fig. 2A, we found a significant negative relationship between increasing water temperature and latency length. In choosing a temperature for the assay, it is important to consider how different temperatures may be optimal based on the goal of the study. As the current study focused on distinguishing differences in baseline pain sensitivity, it was important to select a temperature low enough to be capable of showing variability between groups but high enough to reliably induce a robust pain response. Thus, 48 °C modeled an optimal temperature across the strains and sexes tested.

Although it is not commonly reported, water mixing speed is likely another factor contributing to some discrepancies among previous literature. Higher water mixing speeds are concomitant with increased rates of heat exchange which can thereby accelerate the speed at which thermal pain is induced. To test this effect, we performed the tail withdrawal assay on six WMI males at different water mixing speeds (low, medium, and high) at 48 °C ± 0.25. As shown in Fig. 2B, we observed significantly shorter latencies at faster water mixing speeds with the longest latencies revealed when the water was not being mixed (i.e., still). These findings elucidate the importance of controlling water mixing speed throughout subsequent experimentation using the tail immersion assay.

One innovative aspect of *TailTimer* is the use of RFID as the primary user data input device. Rodent behavioral tests are usually conducted in tight spaces in which gloves are mandatory. This unique work environment poses challenges for using a keyboard/mouse combination or touch screens as primary data input devices. We therefore adapted the RFID system, primarily used to record the identity of the animals, for use as the primary user interface which can be operated by scanning unique RFID fobs. This implementation provides a convenient solution for entering predetermined information, such as user ID, selection of temperature, progression to the next step, repetition of the measure, etc. The RFID reader we use has a USB interface and is easy to program. Although a keyboard can still be used with *TailTimer*, we almost always use the RFID system because of its convenience.

It is worth noting that *TailTimer* does not include methods of restraining rats that have been utilized across many extant tail immersion studies. The use of restraining methods during testing has been shown to induce additional stress and consequently affect tail withdrawal latency (Huang & Shyu, 1987; Ramabadran et al., 1989). Therefore, we chose to hold the rats without restraint throughout testing. Furthermore, our design does not include a heating device. Due to its general availability in the lab, we used a standard hot plate to maintain the target water temperature at ± 0.25 °C. Subsequent research may aim to establish an alternative to the hot plate that could provide increased temperature stability and thus further abate the degree of variability in water temperature across tests.

*TailTimer* is one of the many laboratory devices that uses the Raspberry Pi single-board computer. These computers encompass computing power that rivals desktop computers of the last decade at a small fraction of their predecessors’ size and cost. A large part of the appeal of these computers is the vast array of external devices (i.e., sensors or motors) that can be connected and controlled through the GPIO ports. Devices for operant conditioning (Longley et al., 2017; Mazziotti et al., 2020), conditioned place preference (Vassilev et al., 2020), head-fixed mesoscale cortical imaging (Murphy et al., 2020), and even virtual reality (Tadres & Louis, 2020) have been reported. Further, a wide range of environmental factors (e.g., humidity, barometric pressure, and light) can be monitored using Raspberry Pi (Longley et al., 2017). Although some technical know-how is needed for making these devices, detailed instructions are generally available. Most of these devices are open source and can therefore be modified to fit new requirements. For example, it is feasible to use *TailTimer* with some modification to measure rectal temperature of rodents.

## Conclusions

In summary, we report an open source, simple, and inexpensive device for measuring the tail withdrawal latency of rodents using the tail immersion assay. This device automatically records latencies, water temperature, and the identifications of both animals and technicians. It also limits the tests to be conducted only when the water is within ± 0.25 °C of the target temperature. We anticipate the increased ease of operation and reproducibility of tail withdrawal latency, provided by *TailTimer*, to augment its utility in nociception and addiction research.

## Notes

https://github.com/chen42/openbehavior/tree/master/tailTimer

### Competing Interest Statement

The authors have declared no competing interest.

