## Supplementary figures and images for "*TailTimer:* an open-source device for automating the rodent tail immersion assay"

### Supplemental Figure 1

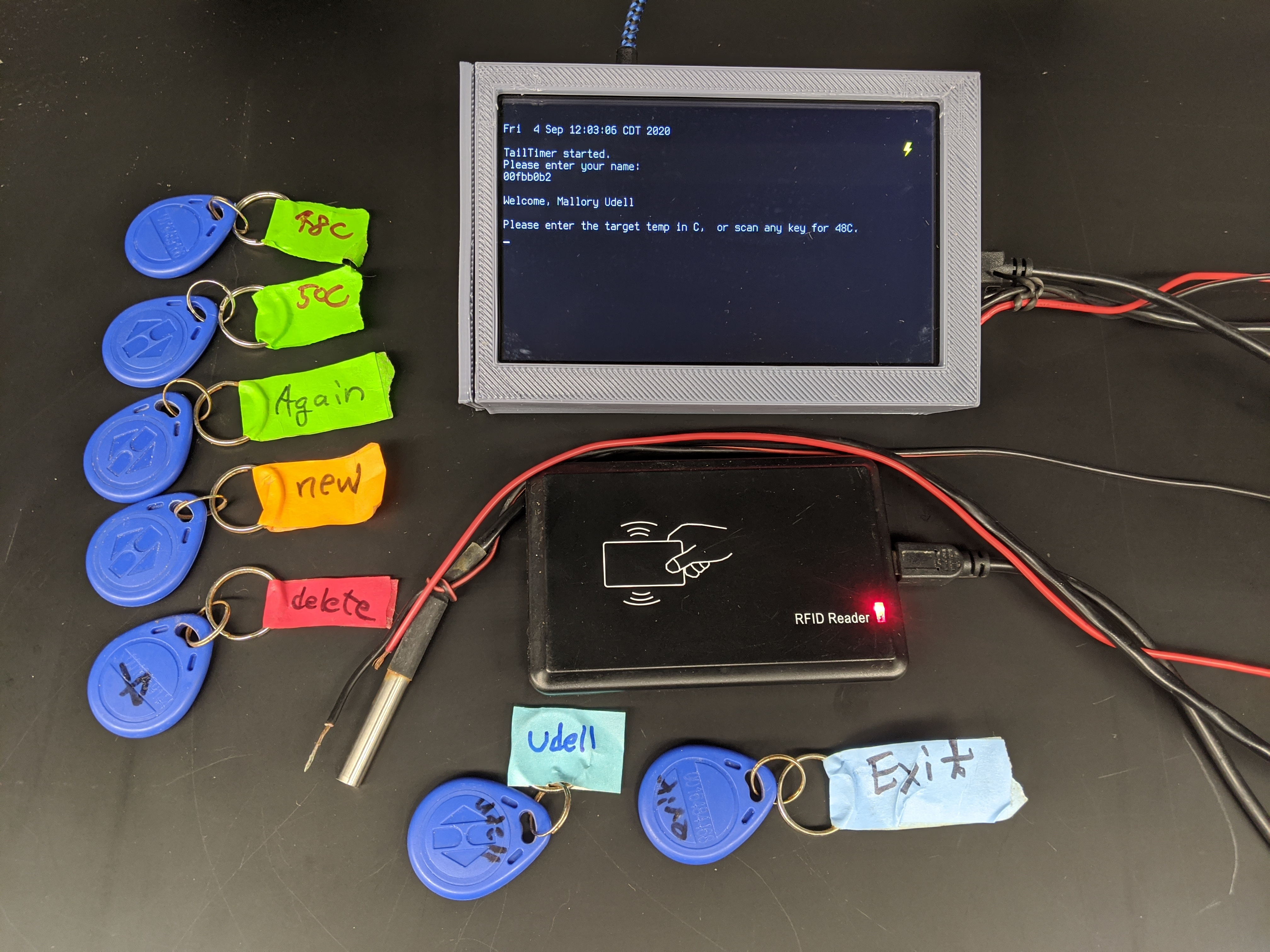
